# Predicting onset, progression, and clinical subtypes of Parkinson disease using machine learning

**DOI:** 10.1101/338913

**Authors:** Faraz Faghri, Sayed Hadi Hashemi, Hampton Leonard, Sonja W. Scholz, Roy H. Campbell, Mike A. Nalls, Andrew B. Singleton

## Abstract

**Background:** The clinical manifestations of Parkinson disease are characterized by heterogeneity in age at onset, disease duration, rate of progression, and constellation of motor versus nonmotor features. Due to these variable presentations, counseling of patients about their individual risks and prognosis is limited. There is an unmet need for predictive tests that facilitate early detection and characterization of distinct disease subtypes as well as improved, individualized predictions of the disease course. The emergence of machine learning to detect hidden patterns in complex, multi-dimensional datasets provides unparalleled opportunities to address this critical need.

**Methods and Findings:** We used unsupervised and supervised machine learning approaches for subtype identification and prediction. We used machine learning methods on comprehensive, longitudinal clinical data from the Parkinson Disease Progression Marker Initiative (PPMI) (n=328 cases) to identify patient subtypes and to predict disease progression. The resulting models were validated in an independent, clinically well-characterized cohort from the Parkinson Disease Biomarker Program (PDBP) (n=112 cases). Our analysis distinguished three distinct disease subtypes with highly predictable progression rates, corresponding to slow, moderate and fast disease progressors. We achieved highly accurate projections of disease progression four years after initial diagnosis with an average Area Under the Curve of 0.93 (95% CI: 0.96 ± 0.01 for PDvec1, 0.87 ± 0.03 for PDvec2, and 0.96 ± 0.02 for PDvec3). We have demonstrated robust replication of these findings in the independent validation cohort.

**Conclusions:** These data-driven results enable clinicians to deconstruct the heterogeneity within their patient cohorts. This knowledge could have immediate implications for clinical trials by improving the detection of significant clinical outcomes that might have been masked by cohort heterogeneity. We anticipate that machine learning models will improve patient counseling, clinical trial design, allocation of healthcare resources and ultimately individualized clinical care.

## Introduction

Parkinson disease (PD) is a complex, age-related neurodegenerative disease that is defined by a combination of core diagnostic features, including bradykinesia, rigidity, tremor and postural instability (Hughes et al. 1992; Postuma et al. 2015). Substantial phenotypic heterogeneity is well recognized within the disease which complicates the design and interpretation of clinical trials and limits counseling of patients about their disease risk and prognosis. The clinical manifestations of PD vary by age at onset, rate of progression, associated treatment complications, as well as the occurrence and constellation of motor/nonmotor features.

The phenotypic heterogeneity that exists within the PD population poses a major challenge for clinical care and clinical trials design. A clinical trial has to be suitably powered to account for interindividual variability, and as a consequence, they are either large, long, and expensive, or only powered to see dramatic effects. This problem becomes particularly burdensome as we move increasingly toward early-stage trials when therapeutic interventions are likely to be most effective. The ability to predict and account for even a proportion of the disease course has the potential to significantly reduce the cost of clinical trials and to increase the ability of such trials to detect treatment effects.

Attempts thus far at characterization of disease subtypes have followed a path of clinical observation based on age at onset or categorization based on the most observable features (Stebbins et al. 2013). Thus, the disease is often separated into early-onset versus late-onset disease, slow progressive “benign” versus fast progressing “malignant” subtypes, PD with or without dementia, or based on prominent clinical signs into a tremor-dominant versus a postural instability with gait disorder subtype (Jankovic et al. 1990; Zetusky, Jankovic, and Pirozzolo 1985). This dichotomous separation, while intuitive, does not faithfully represent the clinical features of disease which are quantitative, complex, and interrelated. A more realistic representation of the disease and disease course requires a transition to a data-driven, multidimensional schema that encapsulates the constellation of interrelated features and affords the ability to track (and ultimately predict) change.

Previous studies used cluster analysis, a data-driven approach, to define two to three clinical Parkinson’s disease subtypes (van Rooden et al. 2010; Fereshtehnejad et al. 2015; Fereshtehnejad et al. 2017). Depth of phenotypic information as well as longitudinal assessments in these studies was variable, and often limited to certain clinical features and short-term follow ups. Moreover, many previous studies were limited by insufficient methods to capture longitudinal changes over multiple assessment visits. A recent study used cluster analysis to identify patient subtypes and their corresponding progression rates (Fereshtehnejad et al. 2017). However, this study evaluated clusters according to only two time points, baseline and short-term follow up, that were aggregated into a Global Composite Outcome score. In return the subtypes did not capture the fluctuations in the prognosis of subtypes. Finally, in order to be used in practice, subtyping solutions need to be replicated in a different cohort and to show the reliability of methods in assigning individual patients to a subtype.

We have previously used multimodal data to produce a highly accurate disease status classification and to distinguish PD-mimic syndromes from PD (Nalls et al. 2015). This effort demonstrated the utility of data-driven approaches in the dissection of complex traits and has also led us to the next logical step in disease prediction: augmenting a prediction of whether a person has or will have PD to include a prediction of the timing and direction of the course of their disease.

Thus, here we describe our work on the delineation and prediction of the clinical velocity of PD. The first stage of this effort requires the creation of a multidimensional space that captures both the features of disease and the progression rate of these features (i.e. velocity). Rather than creating a space based on *a priori* concepts of differential symptoms, we used data dimensionality reduction methods on the complex clinical features observed at 48 months after initial diagnosis to create a meaningful spatial representation of each patient’s status at this time point. After creating this space, we used unsupervised clustering to determine whether there were clear subtypes of disease within this space. This effort delineated three distinct clinical subtypes corresponding to three groups of patients progressing at varying velocities (i.e. slow, moderate and fast progressors). These subtypes were validated and independently replicated. Following the successful creation of disease subtypes within a progression space, we created a baseline predictor that accurately predicted an individual patient’s clinical group membership four years later. This highlights the utility of machine learning as ancillary diagnostic tools to identify disease subtypes and to project individualized progression rates based on model predictions.

## Methods

### Study design and participants

This study included clinical data from the following cohorts: the Parkinson Progression Marker Initiative (PPMI, http://www.ppmi-info.org/; n=328 PD cases including 114 (35%) female; 172 controls including 66 (38%) female), and the Parkinson Disease Biomarkers Program (PDBP, https://pdbp.ninds.nih.gov/; n=112 PD cases including 53 (47%) female; 45 controls including 25 (56%) female). PPMI average age of PD cases was 67±9.8 and control 66.2±11.1. PDBP average age of PD cases 65±8.6 and control 63.7±9.1. The PPMI and PDBP cohorts consist of observational data from comprehensively characterized PD patients and matched controls. All PD patients fulfilled UK Brain Bank Criteria. Control subjects had no clinical signs suggestive of parkinsonism, no evidence of cognitive impairment and no first-degree relative diagnosed with PD. Both cohort’s data went through triage for missing data, 48-month assessment, and comprehensive phenotype collection. Age and MDS-UPDRS Part III (objective motor symptom examination by a trained neurologist) distribution of cohorts at baseline were investigated using Kernel Density Estimation (KDE) to show these independent cohorts are identically distributed and ensure the integrity of replication and validation.

Each contributing study abided by the ethics guidelines set out by their institutional review boards, and all participants gave informed consent for inclusion in both their initial cohorts and subsequent studies.

For each cohort, a comprehensive and shared set of longitudinally collected common data elements were selected for analysis. We used the following data:

i. International Parkinson’s disease and Movement Disorder Society Unified Parkinson’s Disease Rating Scale (MDS-UPDRS) Part-1Part 2, Part 3 (Goetz et al. 2008)
ii. Cranial Nerve Examination
iii. Montreal Cognitive Assessment (Nasreddine et al. 2005)
iv. Hopkins Verbal Learning Test (Brandt 1991)
v. Semantic Fluency test (Goodglass, Kaplan, and Barresi 2001)
vi. WAIS-III Letter-Number Sequencing Test (Wechsler 1997)
vii. Judgment of Line Orientation Test (Benton, Varney, and Hamsher 1978)
viii. Symbol Digit Modalities Test (SMITH and A 1968)
ix. SCOPA-AUT (Visser et al. 2004)
x. State-Trait Anxiety Inventory for Adults (Spielberger et al. 1983)
xi. Geriatric Depression Scale (Yesavage and Sheikh 1986)
xii. Questionnaire for Impulsive-Compulsive Disorders in Parkinson’s Disease (Weintraub et al. 2009)
xiii. REM-Sleep Behavior Disorder Screening Questionnaire (Stiasny-Kolster et al. 2007)
xiv. Epworth Sleepiness Scale (Johns 1991).

### Procedures and statistical analysis

To accompany this report, and to allow independent replication and extension of our work, we have made the code publicly available for use by non-profit academic researchers (https://github.com/ffaghri1/PD-progression-ML). The code is part of the supplemental information; it includes the rendered Python Jupyter notebook with full step-by-step statistical and machine learning analysis. For readability, machine learning parameters have been described in the Python Jupyter notebook and not in the text of the paper. Fig 1 illustrates a summary of our analysis workflow. As a first step, we transformed the dataset into a mathematically meaningful and naturally interpretable format. To achieve this objective, we a) *normalized* and b) *vectorized* all longitudinal data. Specifically, we first vectorized by transforming all observations of a particular parameter in a column vector, then appended all parameters together. We then used the *min-max* method to normalize the data. The min-max method is preferred for multi-modal longitudinal datasets compared to z-score, as it preserves the progression pattern.

**Fig 1.**
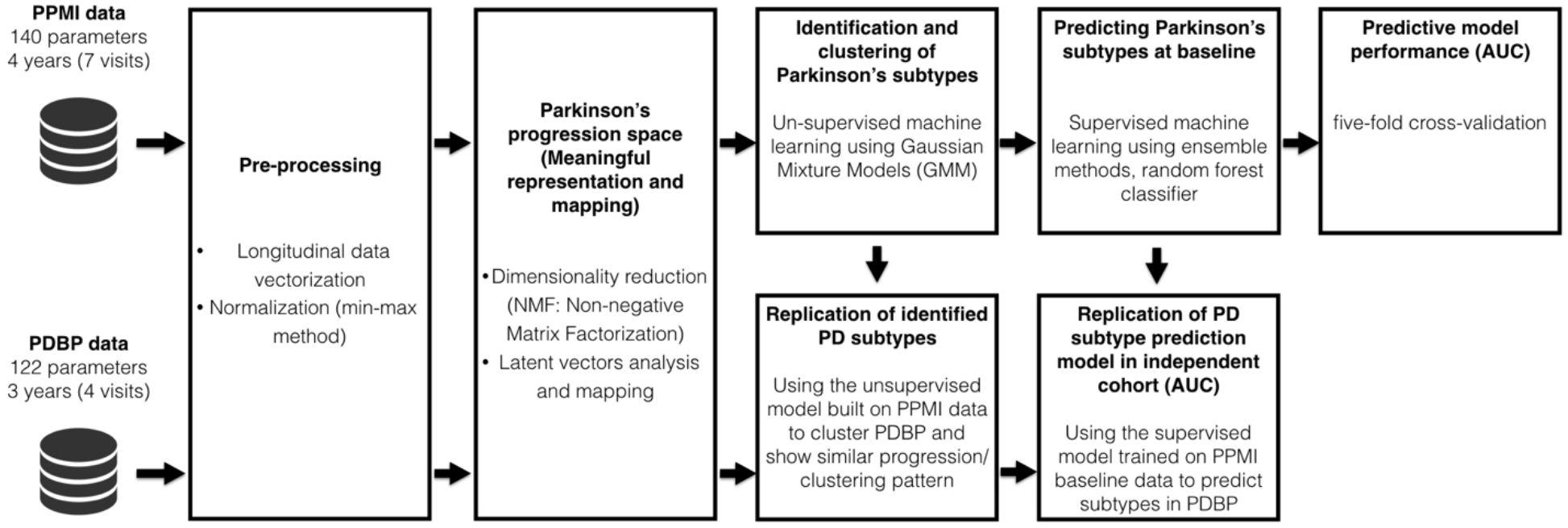
Workflow of analysis and model development.

To develop an interpretable representation of high modality longitudinal data, we next used the dimensionality reduction techniques. Dimensionality reduction techniques helped us to build the “progression space” where we can approximate each patient’s position after 48-month period. We used Non-negative Matrix Factorization (NMF) technique to achieve this aim (Lee and Seung 1999). Alternative methods, such as principal component analysis and independent component analysis, did not perform as well as NMF on clinical longitudinal data. This was expected due to the non-negative nature of our clinical tests. This process essentially collapses mathematically related parameters into the same multi-dimensional space, mapping similar data points closely together.

Mathematically, NMF factorizes (deconstructs) the data into two matrices. The large number of patient parameters are aggregated in a model which represents the underlying progression concept. In this particular use case of NMF, one matrix contains the progression space latent vectors, and the second matrix contains progression stand indicators corresponding to the latent vectors. Latent variables link observation data in the real world to symbolic data in the modeled world. By further looking into the matrix with progression space’s latent vectors, we can identify the mapping and consequently the implications (symbolic dimensions of the modeled progression space).

Through our use of NMF, we identify primary progressive symptomatologies of motor, cognitive and sleep-based disturbances. Following this, unsupervised learning via Gaussian Mixture Models (GMM) (McLachlan and Basford 1988) allowed the data to naturally self-organize into different groups relating to velocity of decline across symptomatologies, from non-PD controls representing normal aging to PD subtypes. GMM is a variant of mixture models, compared to other methods, the parametrization of a GMM allows it to efficiently capture products of variations in natural phenomena where the data is assumed generated from independent and identically distributed mixture of gaussian (normal) distributions. The assumption of normal distribution (and therefore the use of GMM) is often used for population-based cohort phenomenon (Prentice 1986).

We use the Bayesian Information Criterion (BIC) to select the number of PD clusters (subtypes) (Schwarz 1978). The BIC method recovers the true number of components in the asymptotic regime (i.e. much data is available, and we assume that the data was generated i.i.d. from mixture of Gaussian distributions). To replicate the subtype identification, we apply the GMM model developed in the PPMI data to an independent cohort with varying recruitment strategy and design: the PDBP cohort. We show that identified subtypes in the PDBP are similar to the ones in the PPMI in terms of progression.

After identifying progression classes using unsupervised learning, we built predictive models utilizing supervised machine learning via ensemble methods, random forest classifier (Breiman 2001). In preliminary testing, this approach outperformed other methods, such as support vector machines (SVM) and simple lasso-regression models. Besides the predictive performance, we chose random forest (RF) due to the nature of our data and problem: (i) RF is intrinsically suited for multiclass problems, while SVM is intrinsically two-class, (ii) RF works well with a mixture of numerical, categorical, and various scale features, (iii) RF can be used to rank the importance of variables in a classification problem in a natural way which helps the interpretation of clinical results, (iv) RF gives us the probability of belonging to a class, which is very helpful when dealing with individual subject progression prediction. We develop three predictive models to predict patient’s progression class after 48 months based on varying input factors: (a) from baseline factors, and (b) from baseline and first year factors. We also use a feature extraction method, Recursive Feature Elimination (RFE) in order to find significant parameters in our models.

We validate the effectiveness of our predictive models in two ways. First, we validate the algorithm using a five-fold cross-validation. We randomly divided the PPMI dataset into five subsamples, retained a single subsample as the validation data for testing the model, and the remaining four samples used as training data. We repeated the process five times (the folds), with each of the subsamples used exactly once as the validation data. The performance of the algorithm in each fold was measured by the area under the receiver operating curve (AUC) generated by plotting sensitivity vs 1 - specificity. The five results from the folds were averaged to produce a single estimation of performance.

To conclusively validate the algorithm, we also evaluated the performance of the predictive models (trained on the PPMI measurements) on the independent PDBP cohort. We show that the predictive models preserve their high accuracy applied to another dataset.

## Results

Fig 2 shows the result of mathematical projection of PD progression, called *Parkinson disease progression space*. This space shows the relative progression velocity of each patient in 48 months (i.e. speed and direction). The level of progression velocity is broken down to three main dimensions: motor, cognitive and sleep-related disturbances. Across these trajectories, the unsupervised learning analysis reveals and classifies patients into three main subtypes of PD, relating to rates of disease progression: PDvec1, PDvec2, and PDvec3. This shows that we can now map the primary clinical symptomatology and disease progression velocity from diagnosis in Parkinson’s.

**Fig 2.**
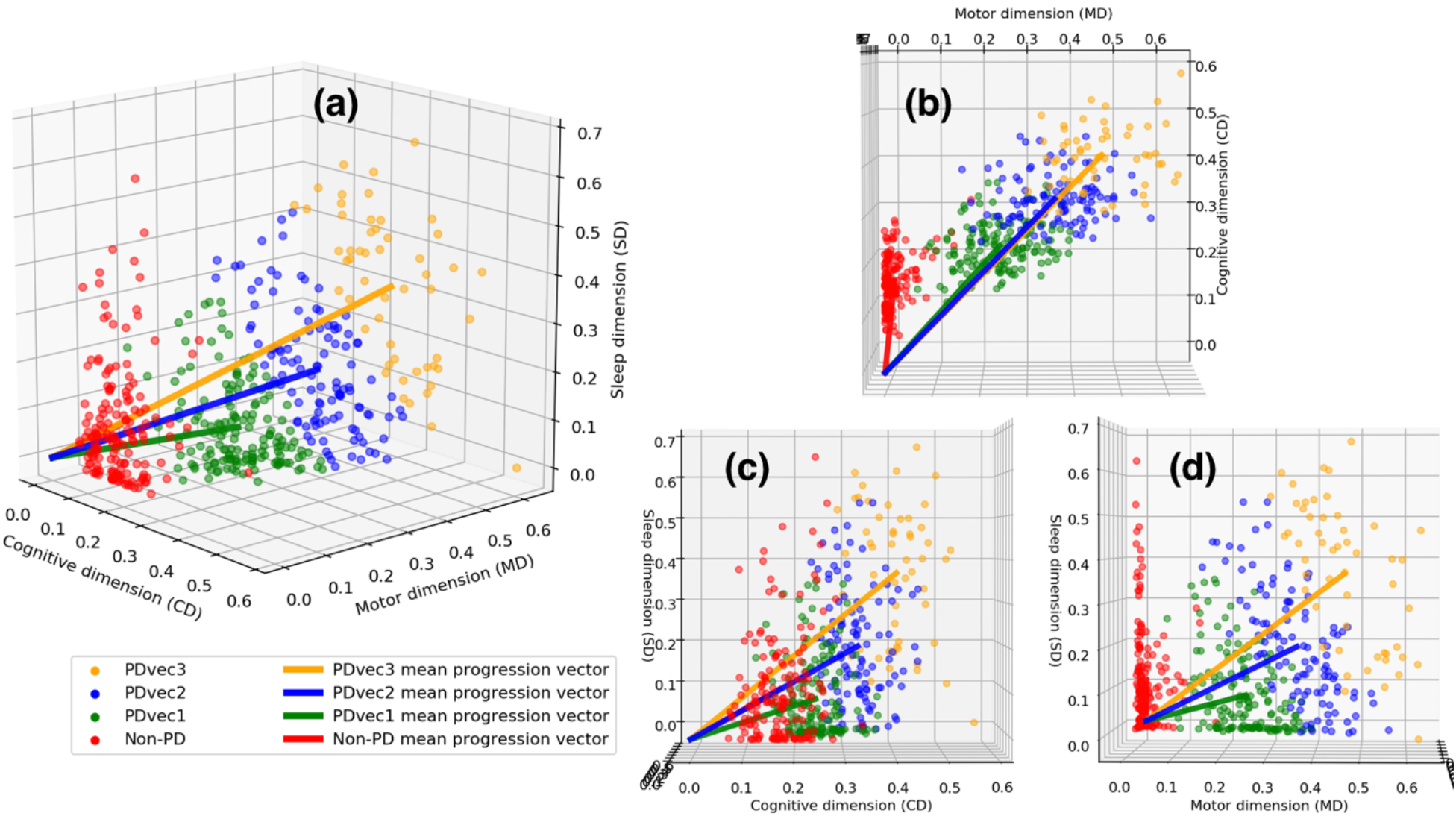
Different views of the Parkinson disease progression space with three corresponding projected dimensions (cognitive, motor, and sleep dimensions). Subtypes of PD are identified using unsupervised learning (PD vector 1, PD vec 2, and PD vec 3). (a) shows the view of all three dimensions, (b) view of motor and cognitive dimensions, (c) view of cognitive and sleep dimensions, and (d) view of sleep and motor dimensions.

Regarding the interpretation of PD progressions space dimensions, Fig 3 shows the mapping guide for how the PPMI’s high-dimensional space of 140 different clinical parameters is mapped to the three-dimensional embedding of Parkinson’s disease progression space. The columns represent the projected three dimensions, i.e. motor, cognitive, and sleep-related trajectories, and the rows are the PPMI clinical parameters. This figure allows us to not only observe the conversion but also the heterogeneity of some clinical parameters, for instance how some of the Epworth Sleepiness Scale parameters reflect both sleep and cognitive disorders and some reflect both sleep and movement disorders. In comparisons of the eigenvalues within the NMF decomposition, the projected motor dimension was responsible for 41·6% of the explained variance, followed by the sleep dimension (29·5%), and cognitive dimension (28·9%).

**Fig 3.**
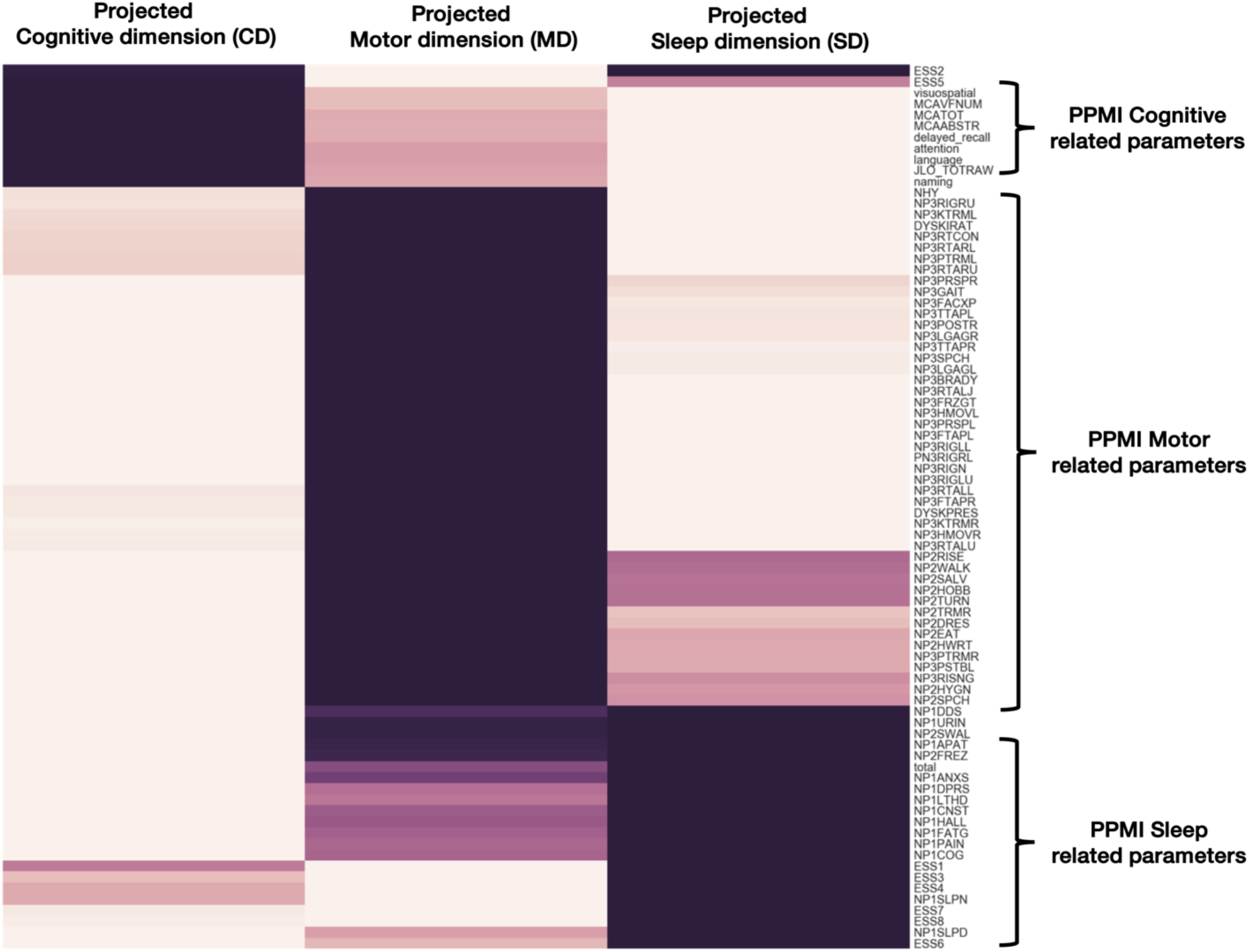
Shows how each 140 different input parameters has been projected into the new dimension of the Parkinson’s progression space (cognitive, motor, and sleep dimensions). Darker colors represent strong mapping.

Regarding the number of identified PD subtypes, Fig 4 shows the characteristics of PDs identified subtypes in more detail. Specifically, Fig 4 shows the visualization of unsupervised learning via GMM. GMM fits the data into different subtypes relating to velocity of decline across symptomatologies, from Non-PD controls. The BIC method has identified three Gaussian distributions representing three PD subtypes.

**Fig 4.**
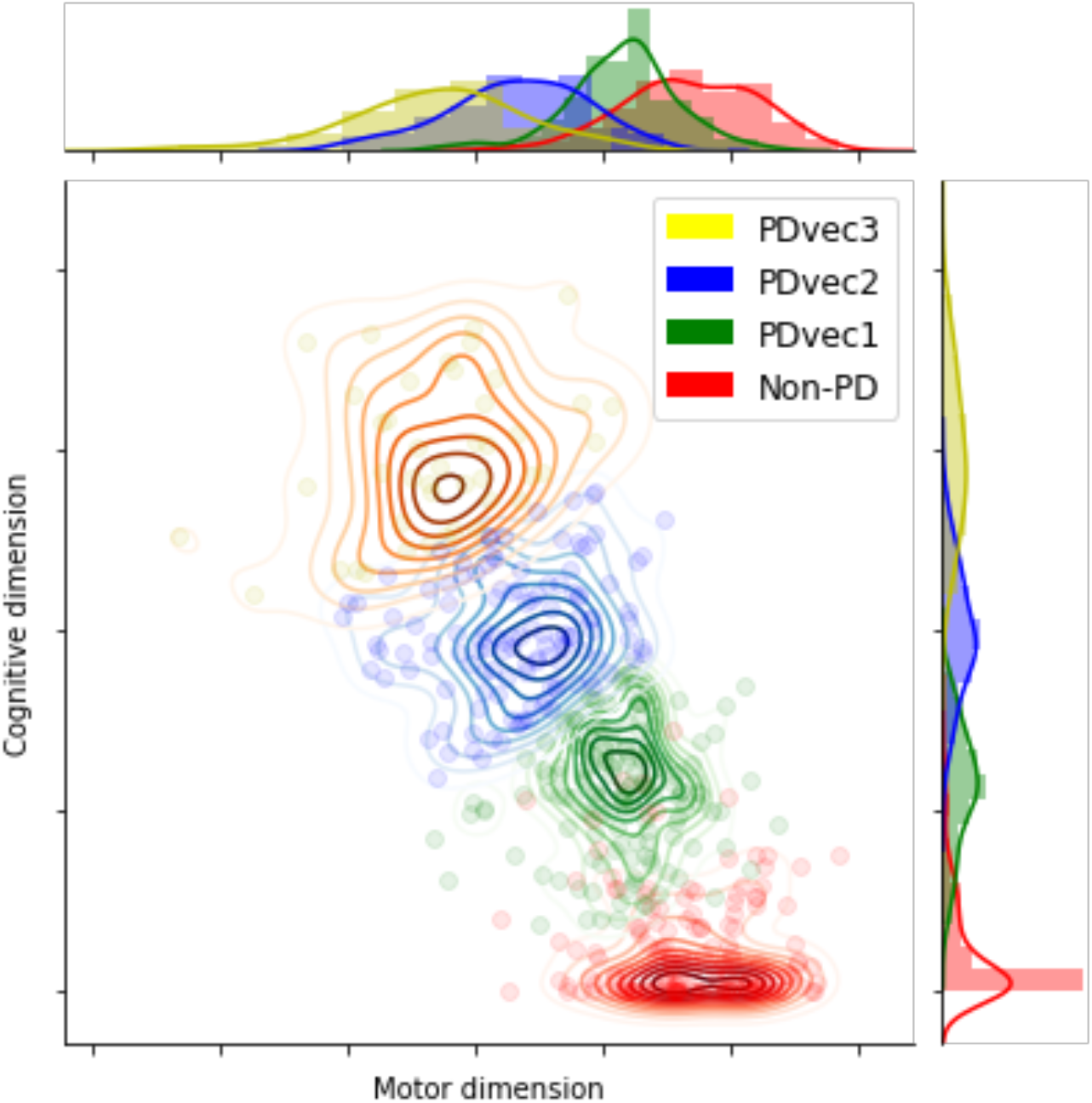
Visualization of unsupervised learning via GMM and identification of three Gaussian distributions representing three distinct PD subgroups. Motor dimension reflects the increase in disturbance, while cognitive dimension reflects the decline.

In terms of characteristics of PDs identified subtypes, Fig 5 demonstrated how cognitive, motor, and sleep-related symptoms differ within each PDs subtype and in controls. There is a clear trend for increased sleep and motor disturbances after four years in fast progressors compared to the slower progressing subtypes which seems to have relatively more cognitive issues early on.

**Fig 5.**
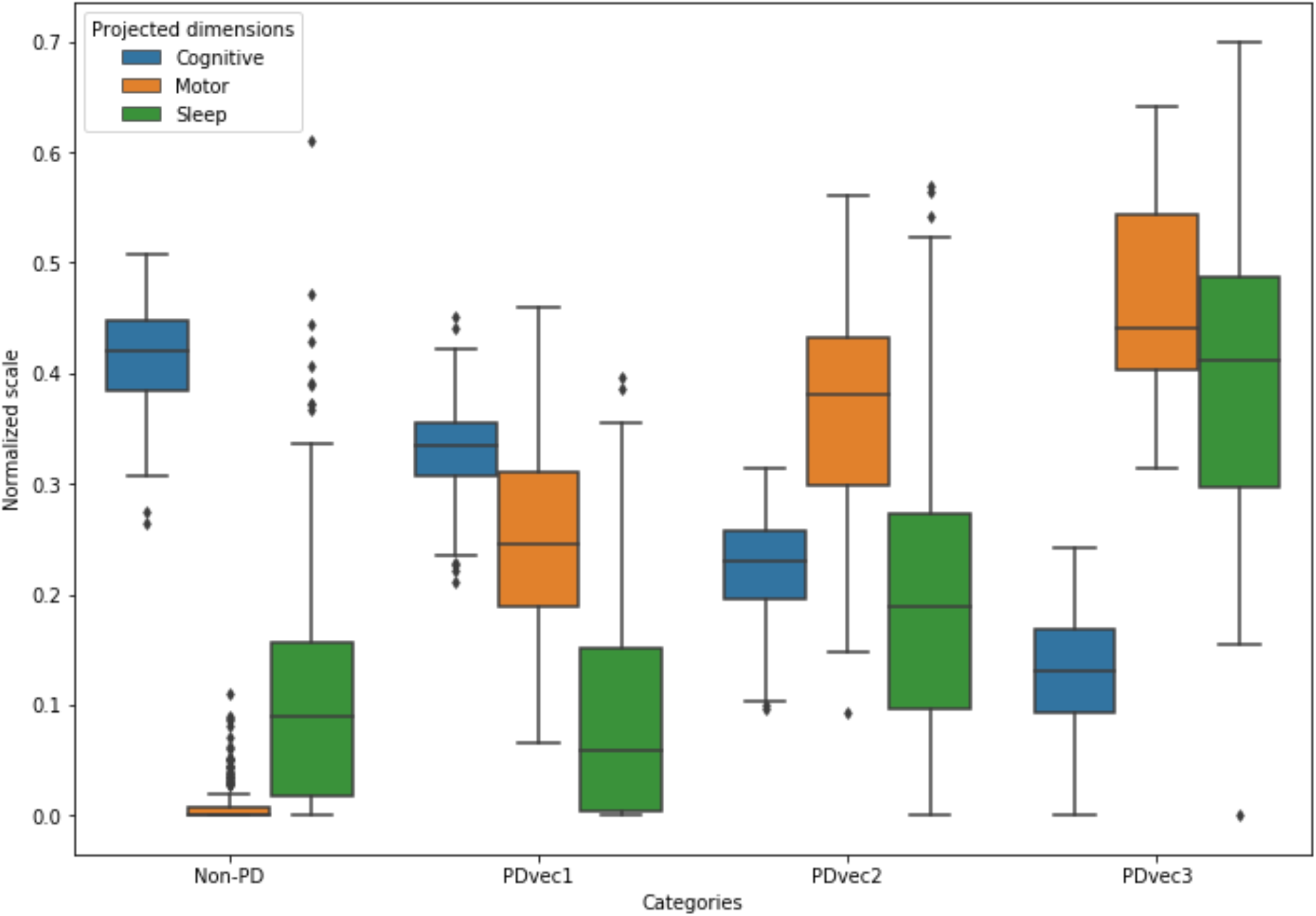
Shows the distribution of projected dimensions (cognitive, motor, and sleep) for each Parkinson category and healthy control after three years. Motor and sleep dimensions reflect the increase in disturbance, while cognitive dimension reflects the decline. PDvec3 has the highest motor and sleep disturbance, as well as highest cognitive decline.

Fig 6 shows the progression of each PD subtype over time at baseline and after 18 months, 24, 36, and 48 months. To better understand the clinical presentation of the three identified subtypes, Fig 6 demonstrates the three main projected dimensions (motor, cognitive and sleep-related disturbances), as well as actual clinical values of each subtype overtime for UPDRS-Part I, Part 2, Part 3, as well as Hopkins Verbal Learning Test, Symbol Digit Modalities Test, Semantic Fluency test, Epworth Sleepiness Scale, State-Trait Anxiety Inventory for Adults, and Geriatric Depression Scale.

**Fig 6.**
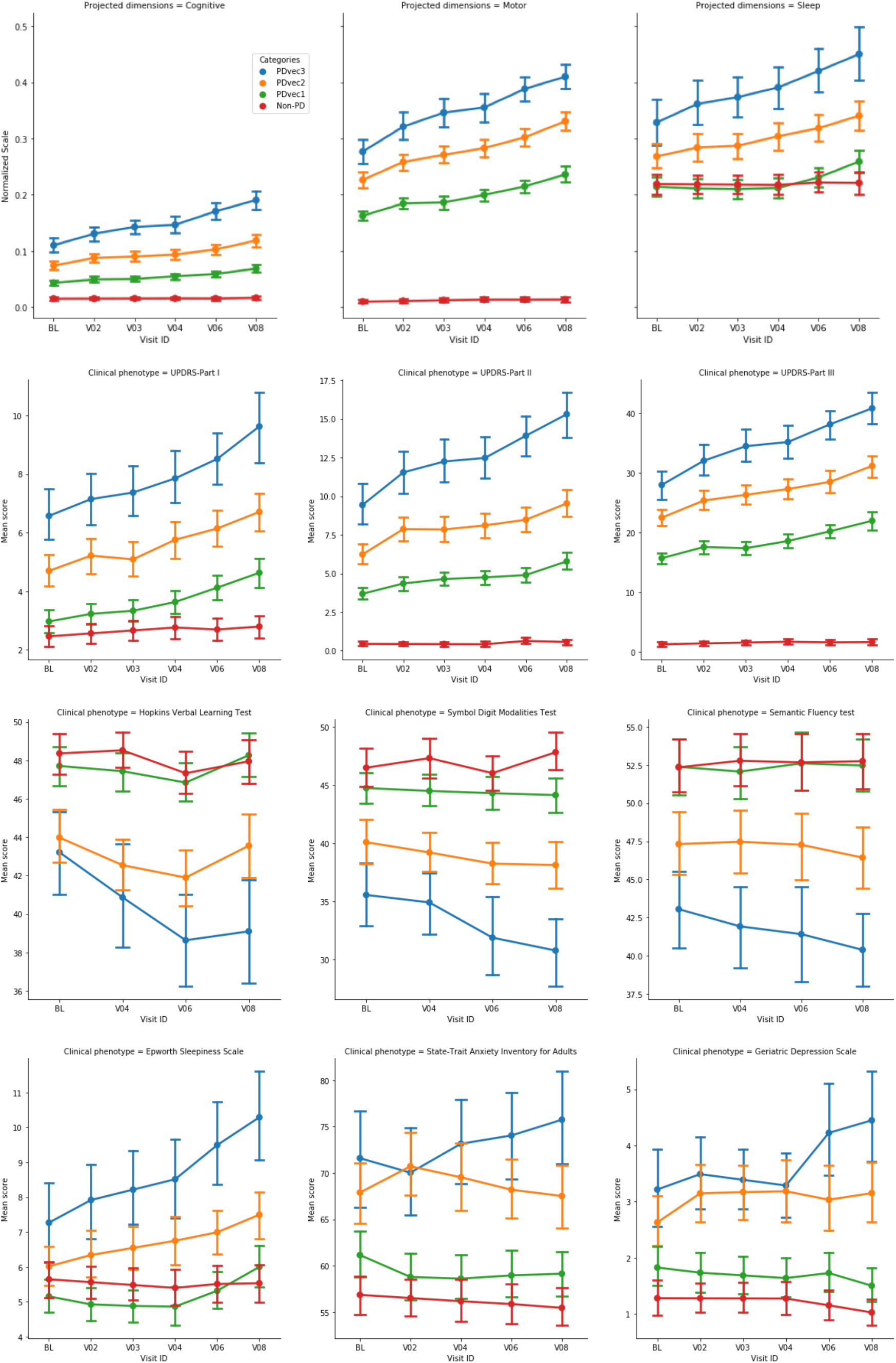
Shows the progression of each PD subgroup over time. Top three graphs show the increased values of motor, sleep, and cognitive dimensions reflect the increase in disturbance overtime. The rest of graphs demonstrate the actual clinical values of each subgroup overtime for UPDRS-Part I, Part 2, Part 3, as well as Hopkins Verbal Learning Test, Symbol Digit Modalities Test, Semantic Fluency test, Epworth Sleepiness Scale, State-Trait Anxiety Inventory for Adults, and Geriatric Depression Scale. BL: Baseline. V03: visit number 3 after 18 months. V04: visit number 4 after 24 months. V06: visit number 6 after 36 months. V08: visit number 8 after 48 months.

In terms of genetic association of PDs identified subtypes, Genetic risk scores (GRS) were calculated as per (Chang et al. 2017). While the GRS was not selected during feature extraction in the clustering exercise we did analyze regressions comparing associations between the GRS and either the continuous predicted cluster membership probability (linear regression) or the binary membership in a particular cluster group compared to the others. All models were adjusted for age at onset, female gender and principal components as covariates to adjust for population substructure in PPMI. The GRS was significantly associated with decreasing magnitude of the sleep vector per Standard deviation of increase in the GRS (beta = −0.0298546, se = 0.0097124, p = 0.00232, adjusted r2 = 0.04584). For binary models of membership, we see that the GRS is weakly but significantly associated with a decreased risk of membership in PDvec3 (odds ratio = 0.5630876 per 1 SD increase from case GRS mean, beta = −0.574320, se = 0.243974, P = 0.01857) and increased risk of membership in PDvec1 (odds ratio = 1.340952, beta = 0.29338, se = 0.13367, P = 0.028178). The lack of strong genetic association is due to the small sample size.

Regarding the replication of subtype identification, Fig 7 shows the identified subtypes in the independent PDBP cohort using the model developed on the PPMI dataset. We see that the identified subtypes in the PDBP are similar to the ones in the PPMI in terms of progression. The PPMI and PDPB cohorts are clinically different cohorts and recruited from different populations. The replication of our results in the PDBP cohort that was recruited with a different protocol shows the strength of our study’s methodology. We demonstrate that if we ascertain the same phenotypes using standardized scales, we can reliably discern the same subtypes and progression rates. This makes our results generalizable and the clinical subtypes reproducible.

**Fig 7.**
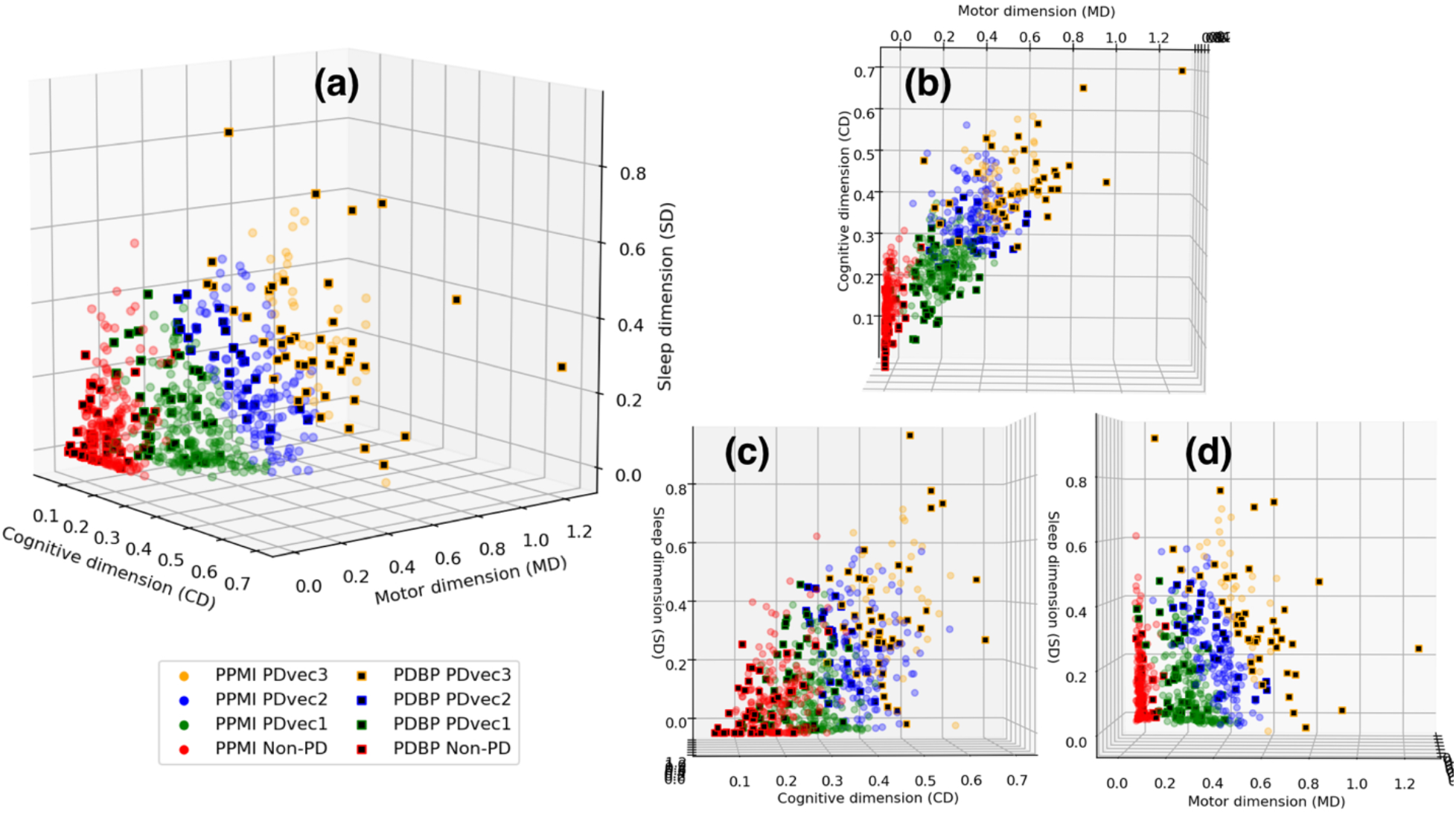
Shows the identified subgroups in the independent PDBP cohort using the model developed on the PPMI dataset. Similar PDBP and PPMI subgroups in terms of progression. (a) shows the view of all three dimensions, (b) view of motor and cognitive dimensions, (c) view of cognitive and sleep dimensions, and (d) view of sleep and motor dimensions.

Following the data-driven organization of subjects into progression subtypes and clustering them into 3 subtypes, we developed three models to predict patient progression class after 48 months based on varying input factors: (a) from baseline factors, and (b) from baseline and year 1 factors. Fig 8a and Fig 8b show the ROC (Receiver Operating Characteristic) curves of our multi-class supervised learning predictors. We correctly distinguish patients with Parkinson’s disease based on baseline only input factors and predict their 48-month prognosis with an average AUC of 0.93 (95% CI: 0.96 ± 0.01 for PDvec1, 0.87 ± 0.03 for PDvec2, and 0.96 ± 0.02 for PDvec3). The predictor built on baseline and year 1 data performs even better with an average AUC of 0.956 (95% CI: 0.99 ± 0.01 for PDvec1, 0.91 ± 0.03 for PDvec2, and 0.97 ± 0.02 for PDvec3). The increased accuracy is due to the availability of more information about a subject. This approach is also practical in a clinical setting, as physicians will provide better prognosis of patients after a year follow up.

**Fig 8.**
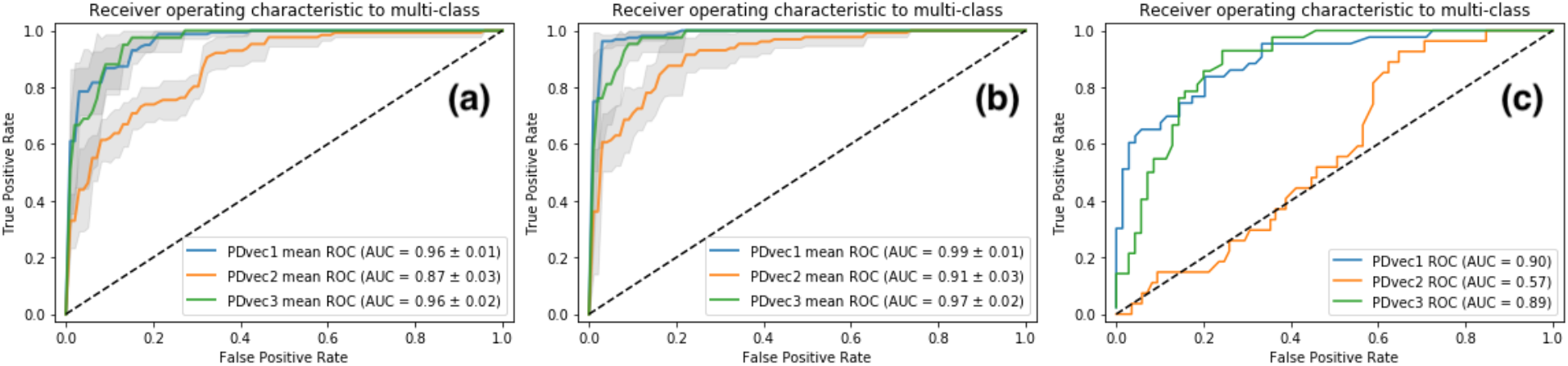
Shows the performance of Parkinson disease progression prediction models. (a) The ROC for the predictive model at baseline developed on the PPMI cohort evaluated using five-fold cross-validation. (b) The ROC for the predictive model developed on baseline and first year data of the PPMI cohort evaluated using five-fold cross-validation. (c) The ROC for the predictive model developed on PPMI baseline and tested on the PDBP cohort.

The predictive model was also analyzed and enhanced by using a feature extraction method: Recursive Feature Elimination. For the predictive model based only on baseline factors, out of 140 clinical parameters, 52 were identified to be the significant contributors (Table 1 for list and detail). Essentially, having only 52 parameters will provide us with highest prediction accuracy. For the predictive model on baseline and year 1 factors, incorporating 66 parameters out of 250 (not all factors were measured at both baseline and first year) provided us with the highest prediction accuracy (Table 1 for list and detail). From these 66 parameters, 34 are baseline measurements and same as baseline predictor, 3 new baseline measurements, and 29 measurements from the first year.

**Table 1.**
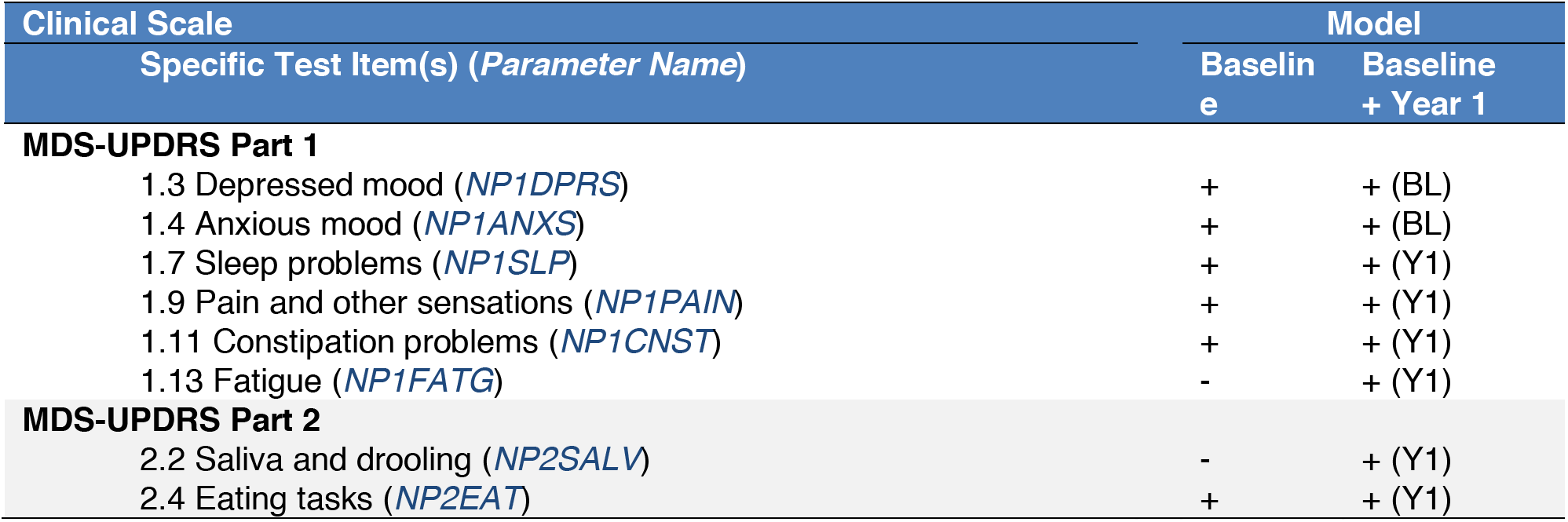
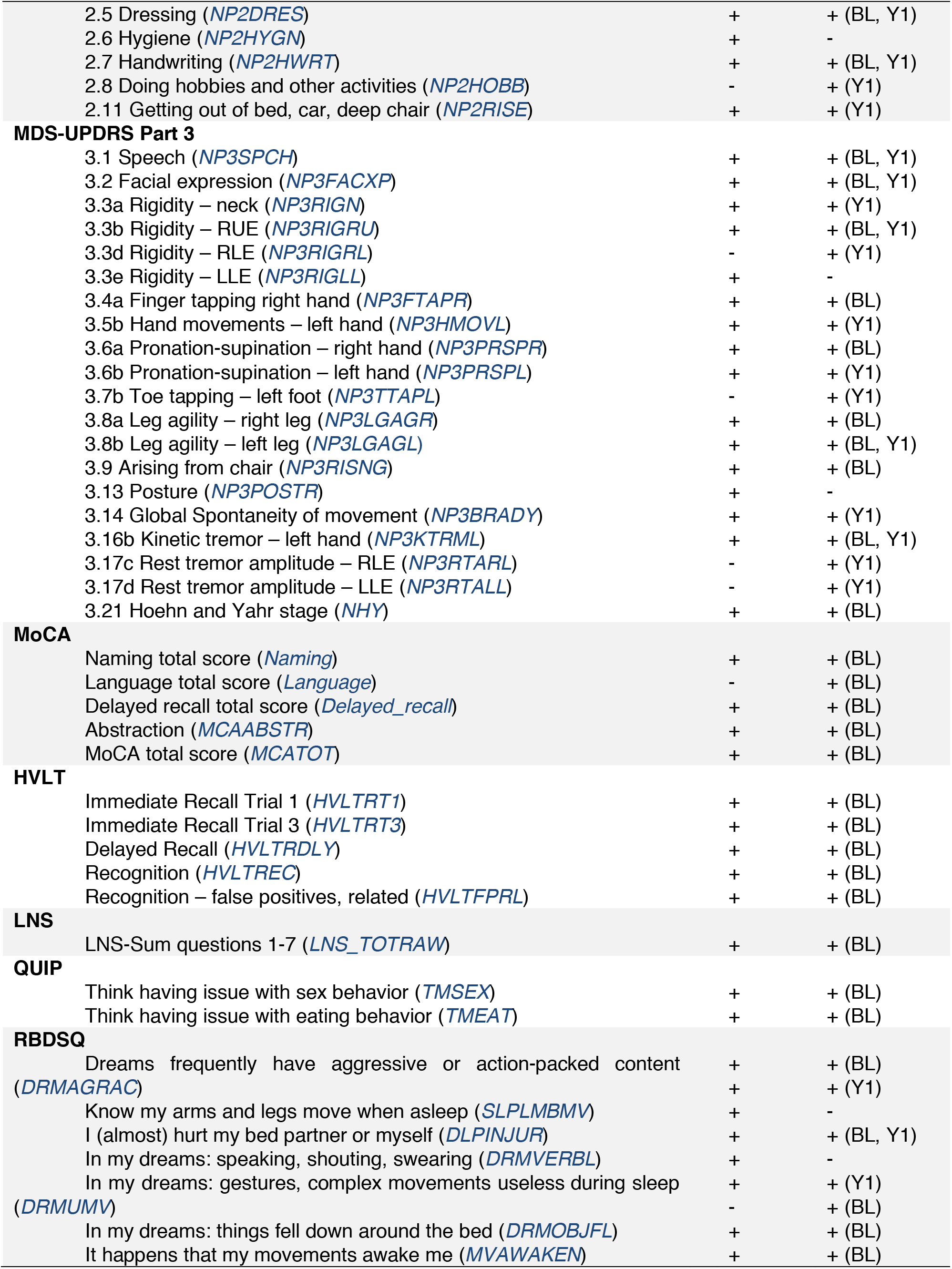
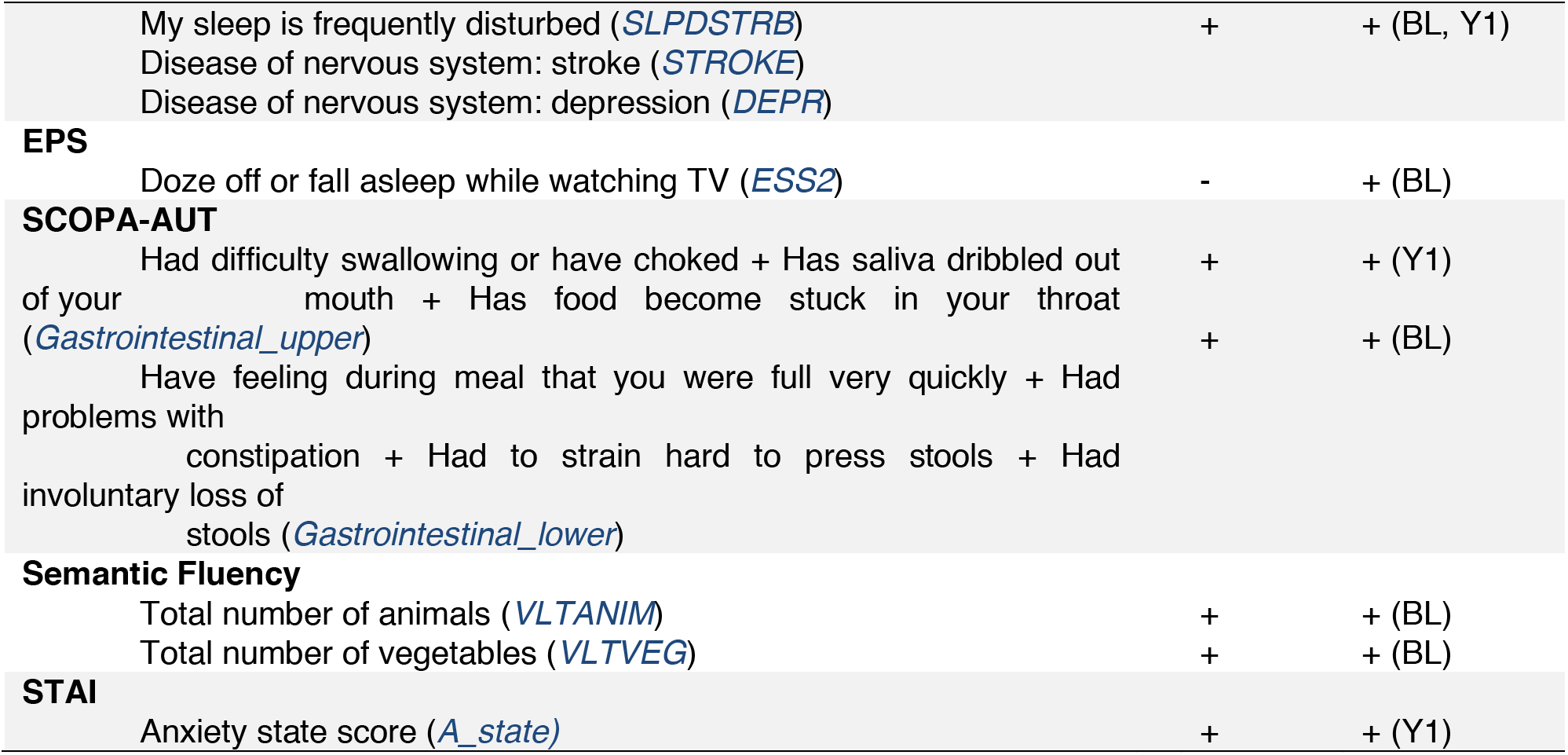
Summary of clinical parameters with significant contributions to the prediction models. Table lists significantly contributing clinical parameters based on baseline examination tests (BL) or based on baseline with year-1 (BL + Y1) test items. Abbreviations: EPS, Epworth Sleepiness Scale; HVLT, Hopkins Verbal Learning Test; LNS, Letter-Number Sequencing; MDS-UPDRS, Movement Disorder Society Revision of the Unified Parkinson’s Disease Rating Scale; MoCA, Montreal Cognitive Assessment; RBDSQ, REM Sleep Behavior Disorder Screening Questionnaire; QUIP, Questionnaire for Impulsive-Compulsive Disorders in Parkinson’s Disease; SCOPA-AUT, Assessment of Autonomic Dysfunction; STAI, State-Trait Anxiety Inventory.

Besides the cross-validation of predictive models in the PPMI cohort, we have also validated the accuracy of predictive model in the independent PDBP cohort. The predictive model trained on the PPMI baseline data correctly distinguished PDBP patients with AUC of 0·787 (ROC curves in Fig 8c). The replicated predictive model performs very well for PDvec1 and PDvec3 (AUC of 0.90 and 0.89 respectively), however, due to small sample size the predictive model does not predict as well on PDvec2 (AUC of 0.57). There are fewer samples that make up the PDvec2 cluster in the replication cohort and it has been easier for the predictive model to predict the more extreme subtypes (i.e. PDvec1 and PDvec3). Despite the smaller sample size of the PDBP cohort, the results strongly validate our previous observations of distinct, computationally discernible subtypes within the PD population. This finding indicates that our methodology is robust, and our unique progression analysis and clustering approach is resulting in the same clusters.

In summary, we have mined data to identify three clinically-related constellations of symptoms naturally occurring within our longitudinal data that summarize PD progression (41.6%, 29.5%, 28.9% variance loadings) comprised of factors relating to motor, sleep and cognitive. We also utilized supervised learning methods to identify the most informative factors across these symptomatologies to predict the velocities of decline for each patient relative to matched health controls with excellent accuracy (>90% after cross validation) from baseline clinical data.

## Discussion

Prediction of disease and disease course is a critical challenge in the care, treatment, and research of complex heterogeneous diseases. Within PD, meeting this challenge would allow appropriate planning for patients and symptom-specific care (for example to mitigate the chance of falls, identifying patients at high risk for cognitive decline or rapid progression etc). Perhaps even more importantly at this time, prediction tools would facilitate more efficient execution of clinical trials. With models predicting a patient-specific disease course, clinical trials could be shorter, smaller, and would be more likely to detect smaller effects; thus, decreasing the cost of phase 3 trials dramatically and essentially reducing the exposure of pharmaceutical companies in a typically expensive and failure-prone area.

We previously had considerable success in constructing, validating, and replicating a model that allows a data-driven diagnosis of PD and the differentiation of PD-mimic disorders, such as those patients who have parkinsonism without evidence of dopaminergic dysfunction (Nalls et al. 2015). We set out to expand this work by attempting to use a somewhat similar approach to 1) define natural subtypes of disease; 2) attempt to predict these subtypes at baseline; and 3) to identify progression rates within each subtype and project progression velocity.

While the work here represents a step forward in our efforts to sub-categorize and predict PD, much more needs to be done. The application of data-driven efforts to complex problems such as this clearly works, however, the primary limitation of such approaches is that they require large datasets to facilitate model construction, validation, and replication. To achieve the most powerful predictions, these datasets should include standardized phenotype collection and recording. Collecting such data is a challenge in PD, with relatively few cohorts available with deep, wide, well-curated data. Thus, a critical need is the expansion or replication of efforts such as PPMI or PDBP, importantly with a model that allows unfettered access to the associated data; the cost associated with this type of data collection is large, but these are an essential resource in our efforts in PD research.

## Acknowledgements

We thank the patients and their families who contributed to this research This work was supported in part by the Intramural Research Program of the National Institute on Aging and the National Institute of Neurological Disorders and Stroke, National Institutes of Health, Department of Health and Human Services (project _ZO1 AG000949, ZIA-NS003154) and the Michael J Fox Foundation. This work has also been supported in part through the grant 1U54GM114838 awarded by NIGMS through funds provided by the trans-NIH big data to Knowledge (BD2K) initiative (www.bd2k.nih.gov). Data used in the preparation of this article were obtained from the Parkinson’s Progression Markers Initiative (PPMI) database (www.ppmi-info.org/data). For up-to-date information on the study, visit www.ppmi-info.org. PPMI – a public-private partnership – is funded by the Michael J. Fox Foundation for Parkinson’s Research and funding partners, including Abbvie, Avid Radiopharmaceuticals, Biogen Idec, Bristol-Myers Squibb, Covance, Eli Lilly & Co., F. Hoffman-La Roche, Ltd., GE Healthcare, Genentech, GlaxoSmithKline, Lundbeck, Merck, MesoScale, Piramal, Pfizer, and UCB. Data and biospecimens used in preparation of this manuscript were obtained from the Parkinson’s Disease Biomarkers Program (PDBP) Consortium, part of the National Institute of Neurological Disorders and Stroke at the National Institutes of Health. Investigators include: Roger Albin, Roy Alcalay, Alberto Ascherio, DuBois Bowman, Alice Chen-Plotkin, Ted Dawson, Richard Dewey, Dwight German, Xuemei Huang, Rachel Saunders-Pullman, Liana Rosenthal, Clemens Scherzer, David Vaillancourt, Vladislav Petyuk, Andy West and Jing Zhang. The PDBP Investigators have not participated in reviewing the data analysis or content of the manuscript.

